# Increased mTOR activity and metabolic efficiency in mouse and human cells containing the African-centric tumor-predisposing p53 variant Pro47Ser

**DOI:** 10.1101/2020.02.14.946269

**Authors:** Keerthana Gnanapradeepan, Subhasree Basu, Thibaut Barnoud, Julia I-Ju Leu, Madeline Good, Joyce V. Lee, William J. Quinn, Che-Pei Kung, Rexford S. Ahima, Joseph A. Baur, Kathryn E. Wellen, Qin Liu, Zachary T. Schug, Donna L. George, Maureen E. Murphy

**Author notes:** Correspondence: Maureen E. Murphy PhD Program in Molecular and Cellular Oncogenesis The Wistar Institute 3601 Spruce St Philadelphia PA 19104.

## Abstract

The Pro47Ser variant of p53 exists in African-descent populations, and is associated with increased cancer risk in humans and mice. This variant, hereafter S47, shows altered regulation of the cystine importer *Slc7a11*, and S47 cells possess increased cysteine and glutathione (GSH) accumulation compared to cells with wild type p53. In this study we show that mice containing the S47 variant have increased mTOR activity, increased oxidative metabolism, larger size, and improved metabolic efficiency. Mechanistically, we show that there is increased association between mTOR and its positive regulator Rheb in S47 cells, due to altered redox state of GAPDH, which normally binds and sequesters Rheb. Compounds that decrease glutathione in S47 cells normalize GAPDH-Rheb complex formation and mTOR activity. The enhanced metabolic efficiency may have been selected for in early Africa, making the S47 variant one of a growing number of cancer-predisposing genetic variants that possesses other positive, potentially selectable attributes.

## Introduction

The p53 tumor suppressor protein serves as a master regulator of the cellular response to intrinsic and extrinsic stress. Mutations in the *TP53* gene occur in more than 50% of human cancers, and this gene is well-known as the most frequently mutated gene in cancer (Hollstein et al., 1991). p53 works to suppress uncontrolled cellular growth and proliferation through various pathways including apoptosis, senescence, cell cycle arrest and ferroptosis (Stockwell et al., 2017; Vousden and Prives, 2009). More recently a role for p53 in the control of metabolism has emerged. The metabolic functions of p53 include the regulation of mitochondrial function, autophagy, cellular redox state, and the control of lipid and carbohydrate metabolism (Berkers et al., 2013; Gnanapradeepan et al., 2018).

As an integral part of its control of metabolism, p53 negatively regulates the activity of mTOR (mammalian target of rapamycin), which is a master regulator of metabolism in the cell. mTOR is a serine-threonine protein kinase that is stimulated by mitogenic signals, and phosphorylates downstream targets that in turn regulate protein synthesis and cell growth (Ben-Sahra and Manning, 2017). mTOR exists in two distinct signaling complexes: mTORC1 is primarily responsible for cell growth and protein synthesis, while mTORC2 plays roles in growth factor signaling, cytoskeletal control and cell spreading (Liu and Sabatini, 2020). Not surprisingly, mTOR activity is frequently upregulated in a diverse range of cancers. Studies have shown that p53 negatively regulates the mTOR pathway through transactivation of several target genes including *PTEN*, *TSC2*, *PRKAB1* and *SESN1/SESN2* (Budanov and Karin, 2008; Feng et al., 2005). The regulation of mTOR by p53 is believed to couple the control of genome integrity with the decision to proliferate (Hasty et al., 2013).

*TP53* harbors several functionally-impactful genetic variants or single nucleotide polymorphisms (SNPs) (Basu et al., 2018; Jennis et al., 2016; Kung et al., 2016). A naturally occurring variant in *TP53* exists at codon 47, encoding serine instead of a proline (Pro47Ser, rs1800371, G/A). This variant exists predominantly in African-descent populations, and occurs in roughly 1% of African Americans and 6% of Africans from sub-Saharan Africa (Murphy et al., 2017). The S47 variant is associated with increased risk for pre-menopausal breast cancer in African American women (Murphy et al., 2017). This variant is defective in the regulation of the small subset of p53 target genes that play roles in ferroptosis sensitivity, including the cystine importer *SLC7A11.* This is accompanied by increased cysteine levels and glutathione (GSH) accumulation in cells from S47 humans and mice (Jennis et al., 2016; Leu et al., 2019). More recently, we showed that the ferroptotic defect in S47 mice leads to iron accumulation in their livers, spleens and macrophages; moreover, the S47 variant is positively associated with iron overload in African Americans (Singh et al., 2020). Of note, the S47 mouse develops significantly increased incidence of spontaneous cancer, particularly liver cancer (Jennis et al., 2016).

An interesting paradigm that is emerging in the literature is that tumor-predisposing genetic variants may paradoxically provide selection benefit to individuals, thus potentially explaining the frequency of these damaging alleles in the population. As an example, women carrying tumor-predisposing mutations in the *BRCA1* gene tend to be physically larger and show increased fertility (Smith et al., 2012). Here-in we show that mice carrying a knock-in S47 allele in a pure C57Bl/6 background show increased size, lean content (muscle), and metabolic efficiency, relative to littermate mice with WT p53. We propose that these attributes may have contributed to a positive selection for this variant in sub-Saharan Africa. Our studies shed further light on the intricate regulation that exists between p53, mTOR activity and metabolic output, in this case mediated by GSH and the control of cellular redox state.

## Results

### Higher basal mTOR activity in cells containing the S47 variant

We previously showed that human lymphoblastoid cells (LCLs) that are homozygous for the S47 variant of p53 are impaired for the transactivation of less than a dozen p53 target genes, compared to LCLs from family members with WT p53 (Jennis et al., 2016). Several of these genes encode proteins that play roles in the negative regulation of mTOR (Budanov and Karin, 2008; Feng et al., 2007). We confirmed via Q-RT-PCR that S47 LCLs show decreased expression of the p53 target genes *SESN1* and *PTEN,* and decreased transactivation of *PRKAB1*, relative to WT cells following cisplatin treatment (Figure 1 - figure supplement 1A and B). These findings prompted us to assess basal mTOR activity in WT and S47 LCLs, and in MEFs from WT and S47 mice. We also analyzed tissues from humanized p53 knock-in (Hupki) mice carrying WT and S47 alleles on a pure C57Bl/6 background, which we previously generated and characterized (Jennis et al., 2016). Western blot analysis of WT and S47 LCLs, and from multiple clones of WT and S47 MEFs, revealed increased p-S6K1 (Thr389) in S47 cells; following normalization to total S6K1, this increase was between 2-3 fold (Figure 1A-B). Because the activity of mTOR is influenced by age and gender, with increased mTOR activity in female and older mice (Baar et al., 2016), we next compared age- and sex-matched pairs of lung and muscle tissue from WT and S47 mice. We found increased levels of p-S6K1 and p-mTOR (Ser2448) in S47 lung and skeletal muscle, relative to WT tissues (Figure 1 C-D). As expected, female mice appeared to have overall higher mTOR activity (Baar et al., 2016); however the trend remains in which S47 mice have increased mTOR activity compared to WT mice (Figure 1C, lane 3-4). Immunohistochemical analysis of these tissues from mice supported these conclusions (Figure 1E). Increased mTOR activity was not seen in all tissues of the S47 mouse, however, and lung and skeletal muscle were the most consistently different between WT and S47 (Figure 1 - figure supplement 1C). We also found that p-S6K1, p-mTOR and p-S6 were the most consistently different read-outs of mTOR activity between WT and S47 cells and tissues; in contrast, we saw limited differences in the level of p-4EBP1 between WT and S47 cells (K. Gnanapradeepan, unpublished data). We did not detect significant differences in p-AKT (Ser473) in WT and S47 cells, suggesting that mTORC1 and not mTORC2 activity is predominantly responsible for the observed differences in mTOR activity (Figure 1 - figure supplement 1D).

**Figure 1.**
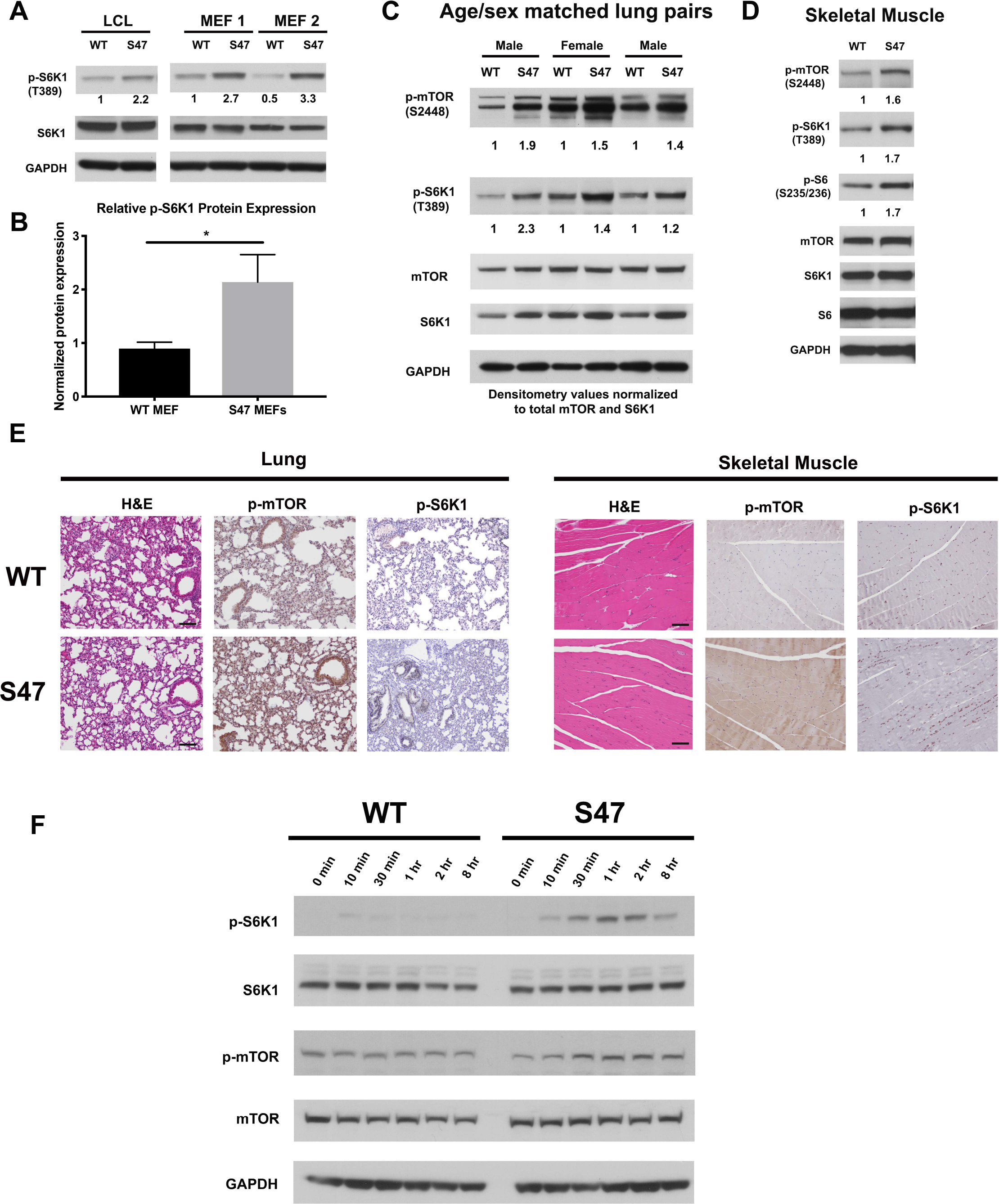
Increased markers of mTOR activity in S47 cells and tissues. (A) Western blot analyses reveal higher phospho-S6K1 expression in S47 LCLs and S47 MEFs, obtained from two separate embryos per genotype. (B) Densitometry quantification of phospho-S6K1 protein expression in WT and S47 MEFs from 4 independent experiments; all values normalized to total S6K1. Error bars represent standard error, (*) p value < 0.05. (C) Whole cell lysates were extracted from 3 WT and 3 S47 mouse lungs and analyzed by Western blot for the proteins indicated. Pair 1 and 3 are lungs isolated from male mice, pair 2 are lungs isolated from female mice. Densitometry quantification of phospho-S6K1 and phospho-mTOR was performed and normalized to total S6K1 and total mTOR protein expression, respectively. (D) Whole cell lysates were extracted from WT and S47 mouse skeletal muscle and analyzed as described above. Densitometry quantification of phospho-S6, phospho-S6K1, phospho-mTOR was performed and normalized to total S6, total S6K1 and total mTOR protein expression, respectively. (E) Immunohistochemical analysis of H&E, phospho-mTOR and phospho-S6K1 in WT and S47 mouse lung and skeletal tissue. Data are representative of n = 4 fields per genotype. Scale bar represents 100 µM. (F) WT and S47 MEFs were starved in media containing 0.1% FBS for 16 hours, then media containing 10% serum was re-introduced and samples were collected at indicated time points. Cell lysates were extracted from samples and subjected to Western Blot analysis for the proteins indicated.

We next sought to challenge mTOR function in WT and S47 cells by subjecting early passage WT and S47 MEFs to a starvation/re-feed experiment, in which we monitored the kinetics of mTOR activation after re-feed using antisera to p-S6K1 and p-mTOR. To do this, three independent cultures each of WT and S47 MEFs were serum starved for 16 hours and refed with 10% serum, after which total and phospho -S6K1 and -mTOR were monitored in a time course. This experiment revealed that S47 cells consistently showed increased induction of markers of mTOR activation after serum re-feed, compared to WT cells (Figure 1F). These data supported the conclusion that the regulation of mTOR activity is distinct in WT and S47 cells.

Given that mTOR plays a role in autophagy inhibition (Jung et al., 2010; White et al., 2011), we hypothesized that basal autophagy, or autophagic flux, might be decreased in S47 cells. However, we were unable to see any differences in the steady state levels of LC3B or the autophagy adaptor protein p62^SQSTM1^ in WT and S47 MEFs or tissues, either at steady state (Figure 1 - figure supplement 1E and F) or following HBSS treatment to induce autophagy (Figure 1 - figure supplement 1F). Likewise, we failed to see differences in viability after HBSS treatment (Figure 1 - figure supplement 1G), nor did we note differences in autophagic flux (conversion of LC3-I to LC3-II when the lysosome is inhibited) in WT and S47 cells (Figure 1 - figure supplement 1H). These data indicate that while markers of mTOR activity are increased in S47 cells and tissues, this does not appear to be accompanied by an inhibition of autophagy.

### Enhanced mitochondrial function and glycolysis in S47 cells

To determine the functional consequences of the increased markers of mTOR activity in S47 cells, we used a Seahorse BioAnalyzer to assess the oxygen consumption rate (OCR), as well as basal and stressed glycolytic rate. Seahorse analyses revealed that S47 LCLs consistently show increased OCR under stressed conditions (Figure 2A). We also found that both human and mouse S47 cells show increased basal and compensatory glycolysis, compared to WT cells (Figure 2B and C). We next sought to more carefully analyze the rates of metabolism in WT and S47 cells by assessing glucose and glutamine consumption using a Yellow Springs Instrument (YSI) Analyzer. We also performed metabolic flux analyses in WT and S47 cells using ^13^C-labeled glucose. Analysis of glucose and glutamine consumption in WT and S47 LCLs and MEFs using the YSI Analyzer revealed that S47 cells show increased consumption of glucose and glutamine, along with increased production of lactate and glutamate, compared to WT cells (Figure 2D-E). We found no evidence that S47 cells proliferate more quickly than WT cells, and if anything S47 LCLs proliferate somewhat less rapidly than WT counterparts (Jennis et al., 2016), suggesting that this increased consumption is being used for biomass instead of proliferation. Interestingly, we also analyzed LCLs from individuals heterozygous for the S47 variant (S47/WT), as well as MEFs from mice heterozygous for this variant, and we noted that these values were typically intermediate between homozygous WT and S47 cells (Figure 2D-E). Analysis of ^13^C6-glucose tracing in WT and S47 MEFs provided evidence for a higher flux of glucose carbon into the TCA cycle in S47 cells compared to WT cells, as evidenced by increased labeling of citrate and malate (Figure 2F-1. G) and aspartate and glutamate (Figure 2 - figure supplement 1A-B). One possibility was that these metabolic changes might be due to increased mitochondrial content, which is regulated by mTOR (Morita et al., 2013). However, MitoTracker analyses and western blotting for mitochondrial proteins revealed no obvious increase in mitochondrial content in S47 cells (Figure 2 - figure supplement 1C-D). The combined data indicate that there is a consistent increase in steady state respiration and glycolysis, as well as increased metabolic flux into the TCA cycle, in S47 cells compared to WT cells.

**Figure 2.**
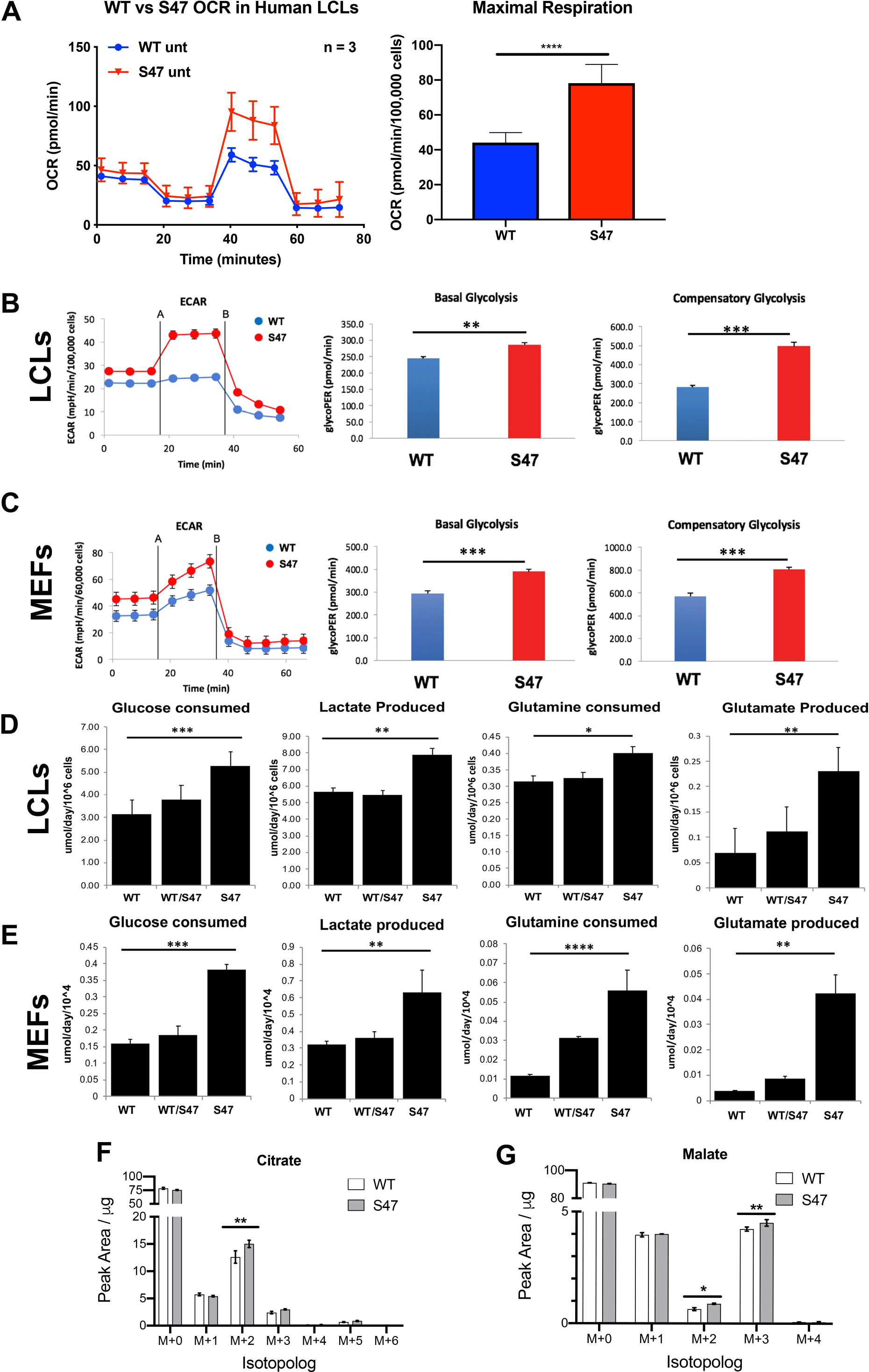
Increased metabolism in S47 cells compared to WT cells. (A) Oxygen consumption rates (OCR) in WT and S47 LCLs were assessed using the Seahorse XF Mito Stress Test. OCR was measured first in basal conditions, then after oligomycin treatment, followed by FCCP treatment and finally rotenone/antimycin. Bar graph depicts maximal OCR after FCCP injection at ∼ 40 minute timepoint; data are representative of 3 independent experiments performed with at least six technical replicates, presented as mean ± SD. (B-C) Basal and compensatory glycolysis in WT and S47 LCLs (B) and MEFs (C) were assessed using the Seahorse Glycolytic Rate Assay. Basal glycolysis is first measured, followed by rotenone/antimycin treatment and finally 2 de-oxy-D-glucose (2-DG) treatment; glycoPER: Glycolytic proton efflux rate. Bar graph depicts basal glycolysis at ∼1 minute timepoint and compensatory glycolysis after antimycin/rotenone injection at ∼ 22 minute timepoint; data are representative of 3 independent experiments performed with at least ten technical replicates. Bar graphs are presented as mean ± SD for all Seahorse analyses. (D-E) Consumption of glucose and glutamine from media and production of lactate and glutamate were analyzed from homozygous WT, heterozygous WT/S47 and homozygous S47 human LCLs (D) and primary MEFs (E) using a YSI-7100 Bioanalyzer. Means and SEM are shown (n= 5). (F-G) WT and S47 MEFs were incubated with 25 mM ^13^C-glucose for 15 minutes and the abundance of citrate (F) and malate (G) isotopologs was quantified by LC-MS/MS. Data are presented as mean ± SD, n =3; 2-way ANOVA. (*) p-value < 0.05, (**) p-value < 0.01, (***) p-value < 0.001, (****) p-value < 0.0001.

The differences in mTOR activity between WT and S47 cells led us to hypothesize that mitochondrial function is altered in these cells, and further that they might respond distinctly to mTOR inhibition, since mTOR is known to regulate mitochondrial metabolism (Morita et al., 2013; Schieke et al., 2006; Ye et al., 2012). Toward testing this hypothesis, we assessed the impact of a non-cytotoxic dose of the mTOR inhibitor, rapamycin, on oxygen consumption rate in WT and S47 cells. Seahorse analysis of WT and S47 LCLs revealed that rapamycin completely abrogates mitochondrial function in WT cells (Figure 3A-C). In contrast, S47 cells are resistant to this dose, particularly when analyzing maximal respiration (Figure 3A-C). These data support the premise that the increased mTOR activity in S47 cells renders their mitochondrial function and oxygen consumption resistant to inhibition of mTOR.

**Figure 3.**
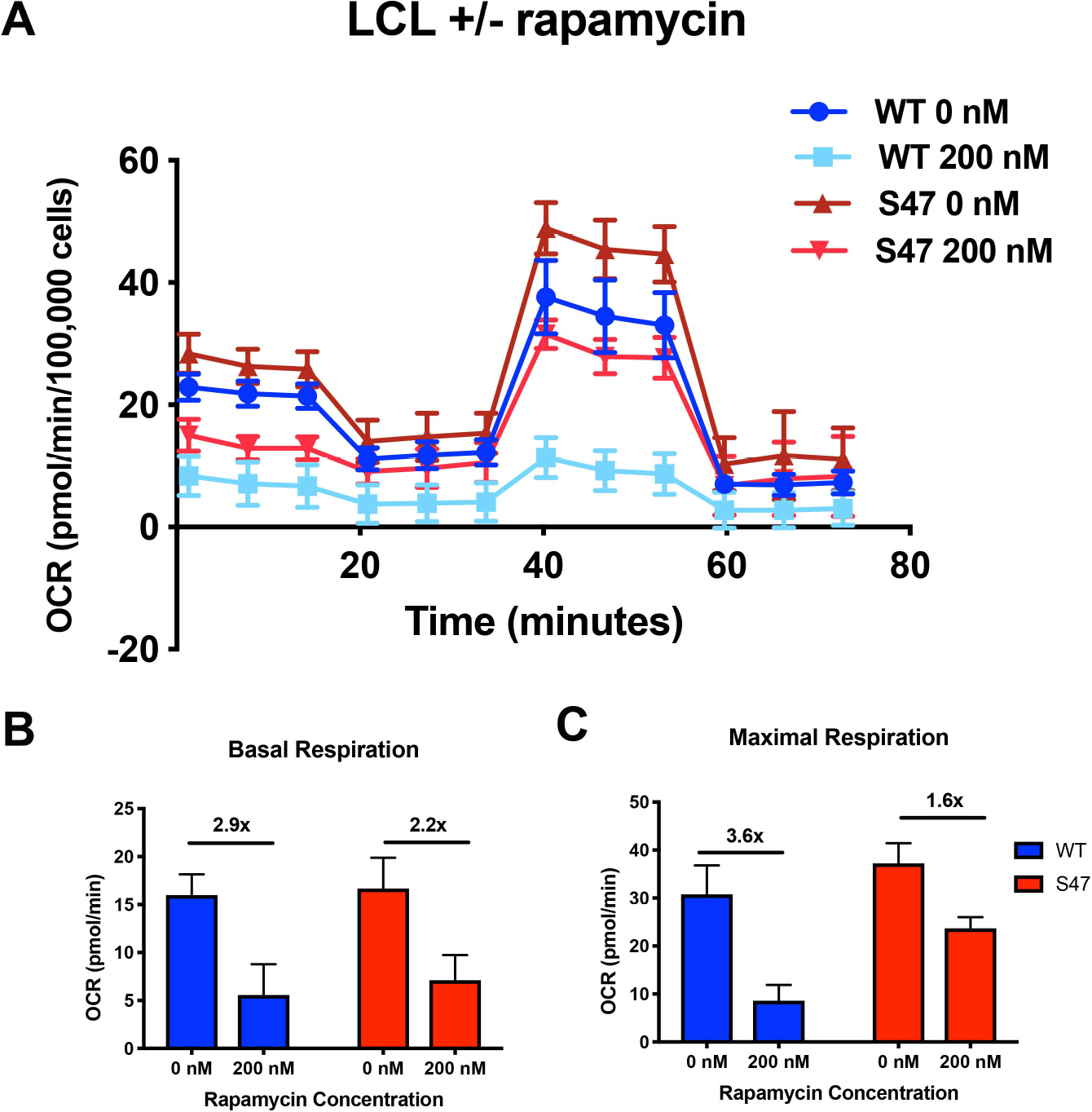
S47 mitochondria show decreased sensitivity to mTOR inhibition. (A) OCR as measured by the Seahorse XFe96 Analyzer in WT and S47 LCLs treated with 200 nM of rapamycin for 24 hours. (B-C) Bar graph depicts basal OCR at ∼1 minute timepoint (B) and maximal OCR after FCCP injection at ∼ 40 minute timepoint (C); fold changes between treated and untreated samples are shown. Data are representative of 2 independent experiments performed with at least ten technical replicates.

mTOR regulates body mass and muscle regeneration (Laplante and Sabatini, 2012; Yoon, 2017). We therefore assessed body weight and composition in age-matched male mice of WT and S47 genotypes. We also tracked body weight with age of multiple male and female sibling littermate mice of WT/WT, WT/S47 and S47/S47 genotypes in our colony. S47 mice showed increased weight with time, compared to WT/WT and WT/S47 sibling littermates (Figure 4 - figure supplement 1A). Body composition analysis using nuclear magnetic resonance revealed that S47 mice had significantly increased fat and lean content, compared to WT mice (Figure 4A-C; Figure 4 - figure supplement 1B-D). We next analyzed WT and S47 mice using a comprehensive lab animal monitoring system (CLAMS) over the course of 48 hours. In this analysis, S47 mice showed reduced food intake, oxygen consumption and heat production, but at the same time comparable locomotor activity (Figure 4D – I). These data support the conclusion that S47 mice possess enhanced metabolic efficiency (less food but equal activity). Before testing this further, we sought to determine the underlying mechanism for increased mTOR activity in S47 cells and mice.

**Figure 4.**
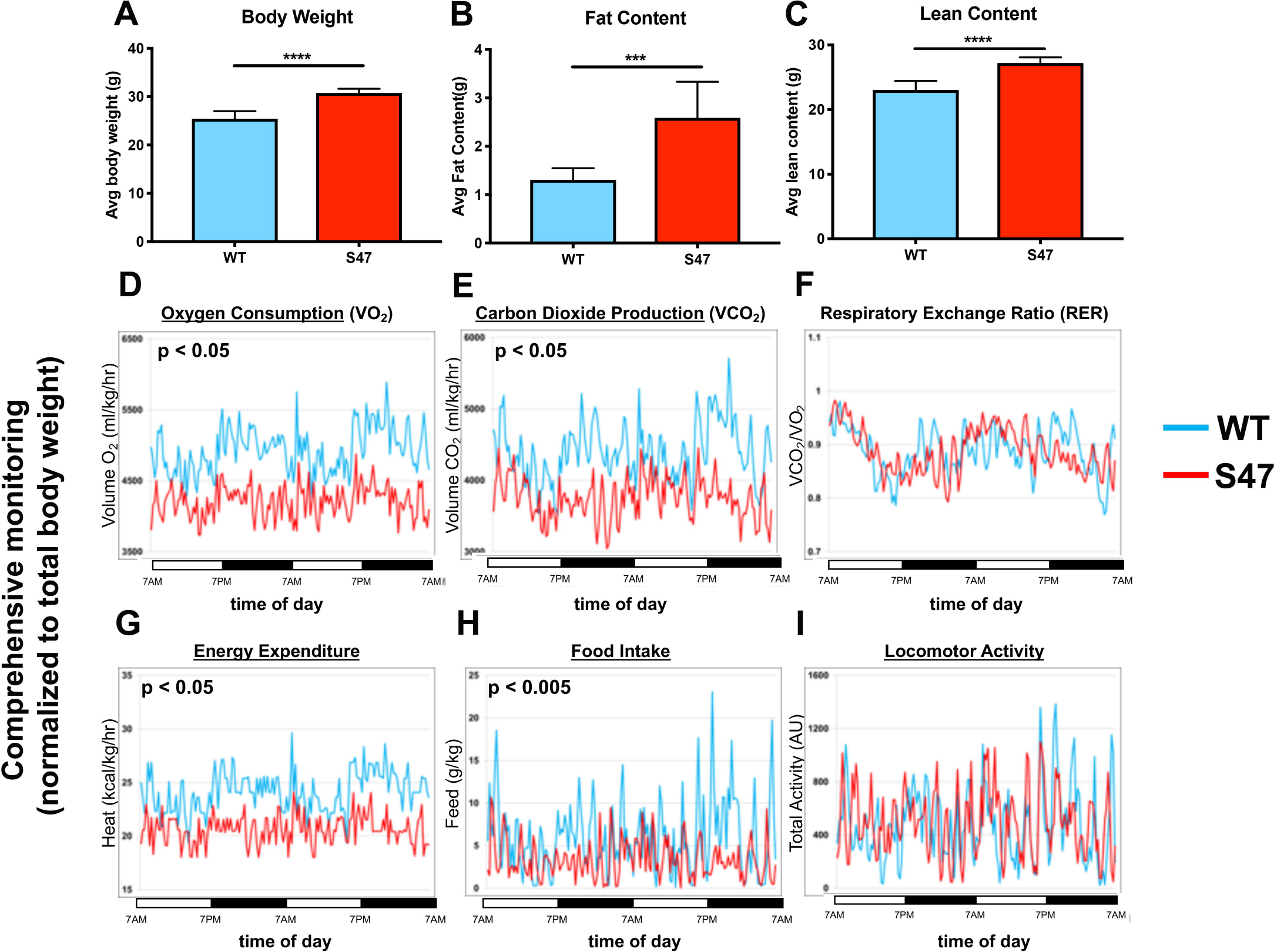
Increased size and improved metabolic efficiency in S47 mice. (A) Nuclear magnetic resonance (NMR) studies revealed S47 mice have increased body weight, (B) increased fat content and (C) increased lean content, n=7 WT mice, n=8 S47 mice. (***) p-value < 0.001, (****) p-value < 0.0001. Bar graphs are presented as mean ± SD. (D-I) Changes in metabolic parameters for WT mice (blue) and S47 mice (red) were determined by using the Comprehensive Lab Animal Monitoring System for 48 hours. Parameters assessed includes (D) oxygen consumption, (E) carbon dioxide production, (F) respiratory exchange rate, (G) energy expenditure, (H) total food intake and (I) locomotor activity. The data are representative of 5 six-week old male mice per genotype and are normalized to total body weight.

### Increased mTOR activity in S47 is due to increased mTOR-Rheb interaction

We were unable to find convincing evidence that the modest differences in gene expression of mTOR negative regulators in S47 cells (Figure 1 – figure supplement 1A) accounted for the differences in mTOR activity between WT and S47 cells. We previously found that S47 cells accumulate iron due to their ferroptotic defect, but we were unable to find evidence that the iron chelator deferoxamine reduced mTOR activity in S47 cells (K. Gnanapradeepan, unpublished observations). Therefore we next analyzed a key regulator of mTOR activity, the small GTPase Rheb, which binds to and activates mTOR (Long et al., 2005). We monitored the mTOR-Rheb association in WT and S47 MEFs using the technique of proximity ligation assay (PLA), which quantitatively detects protein-protein interactions. These PLA experiments indicated that there were consistently increased mTOR-Rheb complexes in S47 cells, compared to WT; this was true in multiple replicates, in multiple MEF clones (Figure 5A-B). One of the recently-identified regulators of the mTOR-Rheb interaction is the cytosolic enzyme GAPDH, which binds to Rheb and sequesters it from mTOR (Lee et al., 2009). Using PLA, we found that the GAPDH-Rheb association is markedly increased in WT cells relative to S47 cells (Figure 5A), despite equivalent levels of all three proteins in cells of both genotypes (Figure 5 - figure supplement 1A). These data indicate that the increased mTOR activity in S47 cells is likely due to increased mTOR-Rheb association, stemming from the decreased GAPDH-Rheb association.

**Figure 5.**
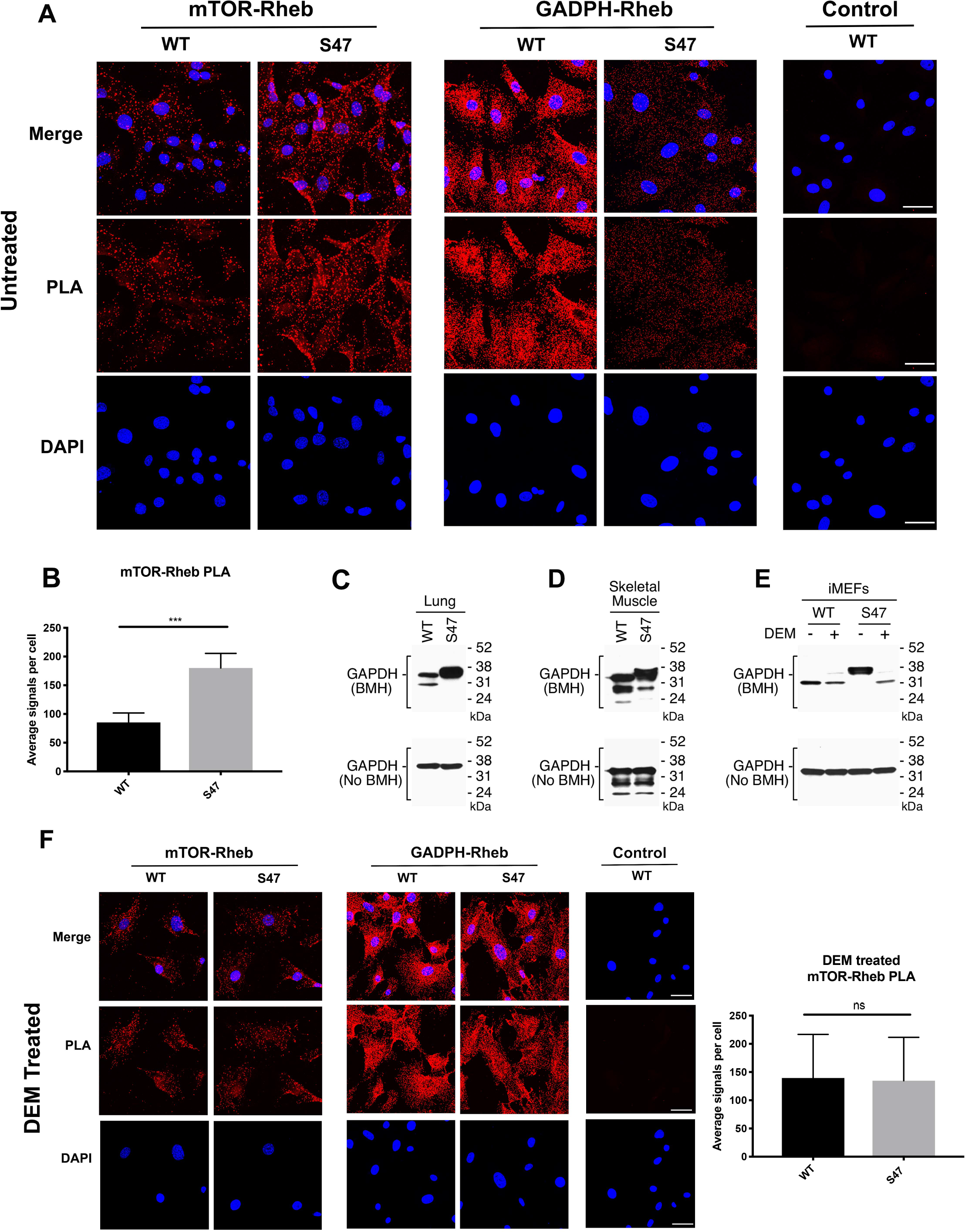
Increased mTOR-Rheb binding in S47 cells, along with altered GAPDH conformation. (A-B) An *in situ* proximity ligation assay (PLA) was performed in WT and S47 MEFs. Each red dot represents an interaction between mTOR-Rheb or GAPDH-Rheb as indicated, scale bar represents 50 µM. The samples were counterstained with DAPI to detect nuclei. Cells stained only with secondary antibody were used as a negative control. (B) Quantification of the mTOR-Rheb interactions, measured as the average number of PLA signals per nuclei. Data were quantified by counting the number of cells in five random fields per experimental condition. (***) p-value < 0.001, Student’s t-test. (C-D) Whole cell lysates were extracted from WT and S47 mouse lung (C) and skeletal (D) tissue. GAPDH proteins were cross-linked with BMH, resolved by SDS/PAGE, and detected by Western blotting with a GAPDH specific antibody (Top). Untreated protein lysates were analyzed by Western blot analysis for GAPDH (Bottom). (E) WT and S47 iMEFs were treated with 50 µM of DEM for 5 h and protein lysates were analyzed as described in *C-D*. (F) PLA was performed in WT and S47 MEFs treated with 50 µM of DEM for 5 h and analyzed as described in *A*.

GAPDH is a multi-functional enzyme that is well-known to be sensitive to redox status (Brandes et al., 2009; Chernorizov et al., 2010). We hypothesized that the increased glutathione (GSH) levels in S47 cells (Leu et al., 2019) might alter the redox state of GAPDH, and likewise alter its ability to bind to Rheb. To assess and compare the redox state of GAPDH in WT and S47 cells, we employed cross-linking experiments using the cysteinyl cross-linking agent bismaleimidohexane (BMH), which cross-links cysteine residues within 13Å by covalently conjugating free (reduced) sulfhydryl groups (Green et al., 2001). BMH was used to treat freshly isolated lung and skeletal muscle lysates from WT and S47 mice, as well as lysates from immortalized WT and S47 MEFs. Cysteinyl-crosslinked proteins were resolved on SDS-PAGE gels, and compared to untreated extracts. We found consistent differences in GAPDH cross-linking patterns in S47 samples compared to WT, as evident by altered mobility on SDS-PAGE of BMH-treated samples (Figure 5C-E). These data suggest that free sulfhydryls are different in WT and S47 cells and tissues, possibly due to the increased GSH in S47 cells. To test this premise further we pre-treated WT and S47 MEFs with the compound diethylmaleate (DEM), which decreases the level of reduced GSH in cells (Leu et al., 2019). We found that pre-treatment of cells with DEM restores the mobility of GAPDH in S47 cells to that evident in cells with WT p53 (Figure 5E). Likewise, we found that depleting excess GSH in cells using DEM or the compound buthionine sulfoximine (BSO, which decreases glutathione synthesis) completely restores GAPDH-Rheb complex formation, and mTOR-Rheb complex formation, in S47 cells to levels equivalent to WT cells (Figure 5F, Figure 5 - figure supplement 1B). Finally, we found that DEM treatment causes decreased p-S6K1 in cells, supporting the premise that GSH levels can influence mTOR activity (Figure 5 - figure supplement 1C). We analyzed the expression of other key regulators of the mTOR and p53 pathway including TSC2, Deptor, AKT and Sco2, and found no significant differences between WT and S47 cells (Figure 5 - figure supplement 1D). The combined data support the conclusion that the increased GSH pool in S47 cells affects the status of reactive cysteines in GAPDH, and the ability of this protein to bind and sequester Rheb, thereby leading to increased Rheb-mTOR interaction and increased mTOR activity in S47 cells.

### Improved treadmill performance of S47 mice

The CLAMs data supporting increased metabolic efficiency in S47 mice prompted us to challenge WT and S47 mice to treadmill exercise with increasing intensity over time; during this time course, oxygen consumption and serum metabolites were quantified. For this analysis we studied eight age-matched male mice of each genotype during a 50 minute forced exercise at increasing speed and slope. Consistent with our CLAMs experiment, S47 mice start with lower basal VO_2_ and exhibit generally lower VO_2_ for the work being performed until they approach the final, most strenuous point of the exercise where the VO_2_ values in WT and S47 converge (Figure 6A-C). Analysis of serum metabolites and proteins before and after exercise revealed decreased lactate dehydrogenase (LDH) levels in S47 mice, indicating less muscle damage in S47 mice compared to WT (Figure 6D). There were no other differences in serum metabolites between WT and S47 mice (Figure 6 - figure supplement 1A). We also monitored the pre-run soleus and gastrocnemius muscle from WT and S47 mice for the steady state level of proteins involved in mitochondrial metabolism, but found no differences in the steady level of any of these proteins (Figure 6 - figure supplement 1B). Finally, we tested WT and S47 males on a continuous, more strenuous, treadmill run and found that three out of four WT mice failed to complete the run, while three out of four S47 successfully completed this run (Figure 6 - figure supplement 1C). Taken together, the combined data point to increased metabolic efficiency and fitness in S47 mice.

**Figure 6.**
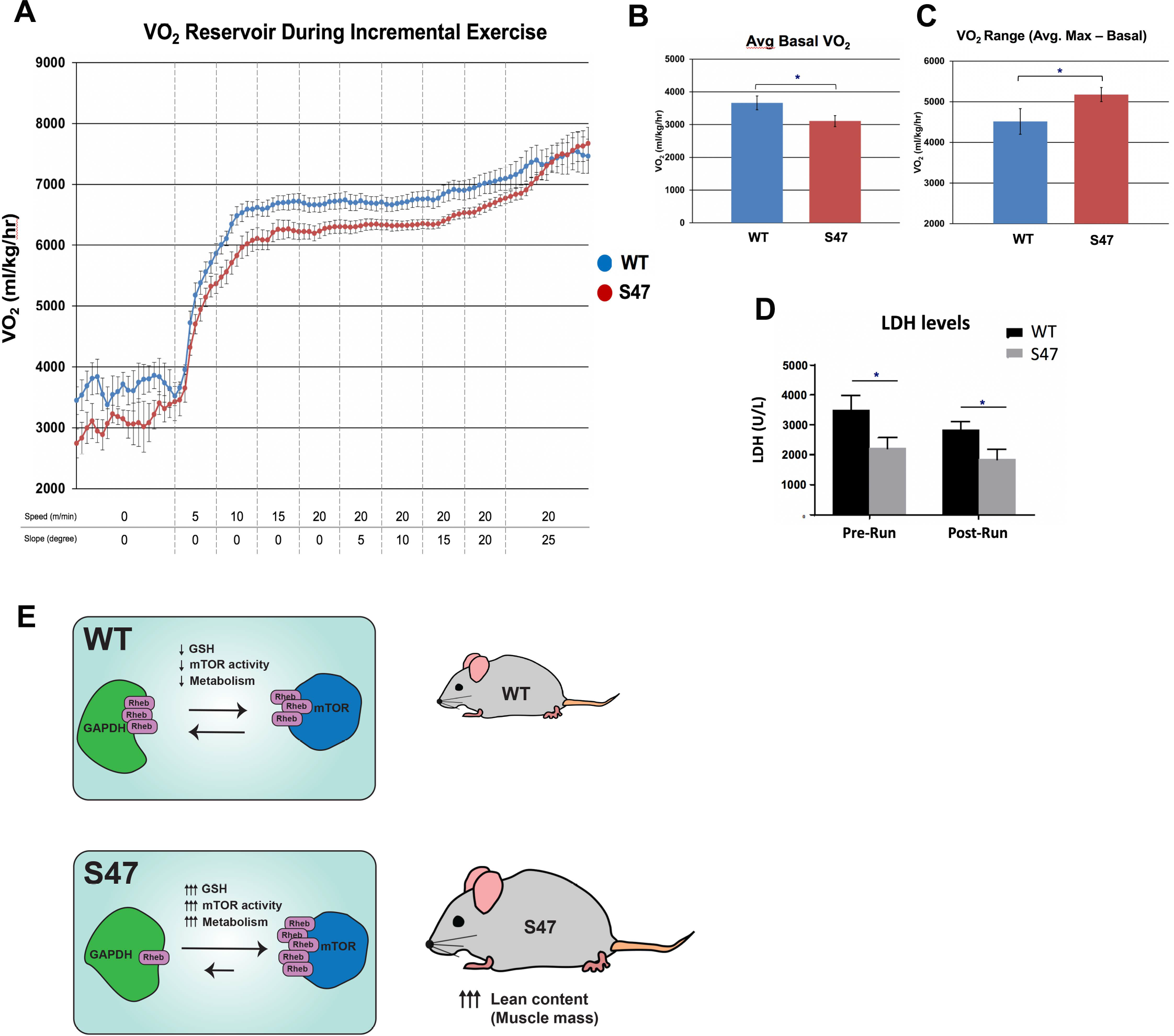
Improved fitness in S47 mice. (A) WT and S47 mice (n = 6-7) were subjected to a treadmill study of increasing intensity over time. Oxygen consumption (VO_2_) is normalized to body mass. (B) Mean basal VO_2_ in WT and S47 mice. (C) VO_2_ range in WT and S47 mice determined by subtracting the mean basal VO_2_ from the VO_2_ max, obtained during the most strenuous point of exercise at the tail end of the treadmill study. (D) Lactate dehydrogenase (LDH) levels measured in the serum of WT and S47 mice obtained before and after the treadmill study. (*) p-value < 0.05, Student’s t-test. (E) Proposed model of how S47 contributes to increased metabolism. The elevated levels of GSH alter the redox state of the S47 cell, in turn affecting GAPDH conformation and impairing GAPDH-Rheb binding. This results in increased mTOR-Rheb binding, leading to increased mTOR activity and resulting an overall increase in metabolism in S47 mice, as seen by increased fat and lean content.

## Discussion

In this study, we show that cells and mice bearing the S47 variant of p53 have increased mTOR activity, increased metabolic efficiency and increased mass. The animals also display signs of superior fitness. The enhanced mTOR activity is due in part to the higher levels of GSH in S47 cells and tissues, which we previously reported (Leu et al., 2019). The increased GSH results in impaired ability of the redox sensitive protein GAPDH to bind to Rheb. This leads to greater mTOR-Rheb binding, resulting in increased mTOR activity in S47 cells and tissues. Our data indicate that, along with pH (Walton et al., 2018), cellular redox status can also regulate mTOR activity, in a manner controlled by p53. We show that oxidative metabolism in S47 cells is less sensitive to mTOR inhibitors, thus tying these two phenotypes together; this is not surprising, as a link between mTOR and a number of cellular metabolic processes is well known (Morita et al., 2013; Schieke et al., 2006). One caveat of this study is that we do not directly demonstrate that the increased mTOR activity in S47 mice is causing their superior performance on treadmill assays; however, heightened mTOR activity is well-known to lead to enhanced muscle recovery after exercise (Song et al., 2017; Yoon, 2017).

We see evidence for increased mTOR activity only in certain tissues of the S47 mouse, so the impact of this genetic variant appears to be somewhat tissue restricted. At present we do not know if this tissue specificity is due to differences in GSH level, or to altered mTOR-Rheb or GAPDH-Rheb interactions in different tissues, or to other parameters. We also see evidence for some unexpected findings regarding the increase in mTOR activity in S47 cells: given that mTOR negatively regulates autophagy (Jung et al., 2010), we expected to see differences in steady state autophagy or autophagic flux in WT and S47 cells, but we found no evidence for this. This finding may be due to the rather complex relationship between mTOR and autophagy (Jung et al., 2010; White et al., 2011), and/or that other signaling pathways regulate autophagy aside from mTOR, including the PI3K pathway, GTPases, and calcium (Yang et al., 2005).

The increased muscle mass in S47 mice likely contributes to the increased fitness observed in these mice. Human studies have shown that mTOR activation is crucial for human muscle protein synthesis, particularly in response to amino acid availability (Dickinson and Rasmussen, 2011). Treatment with the well-studied mTOR inhibitor rapamycin blocks the effects of amino acid ingestion on mTOR activity and leads to decreased protein synthesis in human skeletal muscle (Dickinson and Rasmussen, 2011; Drummond et al., 2009). Additionally, mTOR signaling driven through IGF-1 plays a key role in promoting muscle hypertrophy (Coleman et al., 1995; Musaro et al., 2001; Vandenburgh et al., 1991). Our combined findings suggest that the increased mTOR activity in the muscle of S47 mice leads to the indicators of improved fitness that we see in these mice (see model, Figure 6E). We hypothesize that the more efficient metabolism and enhanced fitness provided by the S47 variant may have once provided carriers with a bio-energetic advantage in Sub-Saharan western Africa, where this variant is most common. For example, those carrying the S47 SNP may have possessed superior athletic prowess and/or ability to withstand famine. This metabolic advantage may explain the high frequency of this genetic variant in sub-Saharan Africa, despite the fact that it predisposes individuals to cancer later in life. A selection for this variant in Africa may also include an improved ability to withstand malaria infection: we recently reported that the S47 variant alters the immune micro-environment in mice, and confers improved response to the malaria toxin hemozoin (Singh et al., 2020). Both of these activities may have conferred selection pressure for this variant in Africa.

Our findings provide further support for the growing premise that some tumor suppressor genetic variants may provide evolutionary selection benefit (Vicens and Posada, 2018). For example, women who carry the BRCA1/2 mutation exhibit increased size and enhanced fertility when compared to controls (Smith et al., 2012). Similarly, people with Li Fraumeni syndrome who inherit germline mutations in *TP53*, as well as mice with tumor-derived germline mutations in *Tp53*, demonstrate increased fitness endurance (Wang et al., 2013), but this is due to increased mitochondrial content, which we do not see in S47 cells. A common genetic variant in *TP53* at codon 72, encoding proline at amino acid 72, confers increased longevity while conversely causing increased cancer risk (Zhao et al., 2018). In contrast, the arginine 72 variant of p53 induces increased expression of LIF, which influences fecundity (Kang et al., 2009). The take home message from all of these studies is that the diverse roles of tumor suppressor proteins like p53 in metabolism, fertility and fitness may allow for positive selection for certain variants, even at the expense of increased cancer risk. In mice, this increased cancer risk occurs quite late in life, well past reproductive selection (12-18 months). More needs to be done to analyze cancer risk in S47 humans. A more comprehensive understanding of the function of tumor suppressor genetic variants, including the S47 SNP, will enable improved understanding of cancer risk, along with superior personalized medicine approaches, with the ultimate goal of improving clinical outcomes and survival of people who carry this variant.

## Materials and Methods

### Mammalian cell culture

WT and S47 MEFs were generated and maintained as previously described (Jennis et al., 2016). Human WT LCLs (Catalog ID GM18870) and S47 LCLs (Catalog ID GM18871) were obtained from the Coriell Institute (Camden, New Jersey) and maintained as previously described (Jennis et al., 2016). MEF cultured cells were grown in DMEM (Corning Cellgro) supplemented with 10% fetal bovine serum (HyClone, GE Healthcare Life Sciences) and 1% penicillin/streptomycin (Corning Cellgro). Human LCLs were grown in RPMI (Corning Cellgro) supplemented with 15% heat inactivated fetal bovine serum (HyClone, GE Healthcare Life Sciences) and 1% penicillin/streptomycin (Corning Cellgro). Cells were grown in a 5% CO_2_ humidified incubator at 37°C. For serum starvation experiments, cells were starved in DMEM containing 0.1% FBS for 16 hours. Following starvation, DMEM containing 10% FBS was re-introduced and cells were harvested at 0 minutes, 10 minutes, 30 minutes, 1 hour, 2 hours and 8 hours after this point. For HBSS experiments, cells were washed once with PBS (Corning 21-031-CV) and then incubated with HBSS (Thermo Fisher Scientific 14025092) for 0, 2 or 6 hours. Viability was assessed using Trypan Blue (Thermo Fisher Scientific 15250061).

### Western blot

For Western blot analyses, 50-100 µg of protein was resolved over SDS-PAGE gels using 10% NuPAGE Bis-Tris precast gels (Life Technologies) and were then transferred onto polyvinylidene difluoride membranes (IPVH00010, pore size: 0.45 mm; Millipore Sigma). Membranes were blocked for 1 hour in 5% bovine albumin serum (Sigma Aldrich, A9647). The following antibodies were used for Western blot analyses: phospho-mTOR 1:1000 (Cell Signaling, 2971), mTOR 1:1000 (7C10, Cell Signaling, 2983), phospho-p70S6K1 1:1000 (Cell Signaling, 9205), p70S6K1 1:1000 (Cell Signaling, 9202), GAPDH 1:10,000 (14C10, Cell Signaling, 2118), TFAM 1:2000 (Abcam, ab131607), MTCO1 1:2000 (Abcam, ab14705), SDHA 1:1000 (Cell Signaling, 5839), Tom20 1:100 (F-10, Santa Cruz, sc17764), phospho-Akt (D9E, Cell Signaling, 4060), p62 1:1000 (Cell Signaling, 5114), LC3B 1:1000 (D11, Cell Signaling, 3868), HSP90 1:1000 (Cell Signaling, 4877S), Rheb 1:1000 (E1G1R, Cell Signaling, 13879), TSC2 1:1000 (D93F12, Cell Signaling, 4308), Akt 1:1000 (Cell Signaling, 9272), Deptor 1:1000 (Novus Bio, NBP1-49674SS). Rabbit or mouse secondary antibodies conjugated to horseradish peroxidase were used at a 1:10,000 dilution (Jackson Immunochemicals), followed by a 5-minute treatment with ECL (Amersham, RPN2232). Protein levels were detected using autoradiography and densitometry analysis of protein content was conducted using ImageJ software (NIH, Rockville, MD).

### Immunohistochemistry

Tissues were harvested and fixed in formalin overnight at 4°C, followed by a wash with 1X PBS and were then placed in 70% ethanol prior to paraffin embedding. The Wistar Institute Histotechnology Facility performed the tissue embedding and sectioning. For the immunohistochemistry (IHC) studies, paraffin embedded tissue sections were de-paraffinized in xylene (Fisher, X5-SK4) and re-hydrated in ethanol (100%-95%-85%-75%) followed by distilled water. Samples underwent antigen retrieval by steaming slides in 10 mM Citrate Buffer (pH 6). Endogenous peroxidase activity was quenched with 3% hydrogen peroxide and slides were incubated in blocking buffer (Vector Laboratories, S-2012) for 1 hr. The slides were incubated with phospho-p70S6K1 (1:100, ThermoFisher Scientific, PA5-37733) or phospho-mTOR (1:100, Cell Signaling, 2971) primary antibody overnight at 4°C. The following day, slides were washed with PBS and incubated with HRP-conjugated secondary antibody for 30 mins. Antibody complexes were detected using DAB chromogen (D5637). Light counterstaining was done with hematoxylin. Slides were imaged using the Nikon 80i upright microscope and at least four fields were taken per section.

### Mitochondrial metabolism assays

The oxygen consumption rate (OCR) and glycolytic rate were determined using the Seahorse XF MitoStress Assay and the Seahorse XF Glycolytic Rate Assay, respectively, according to the manufacturer’s protocol. Cells were plated one day prior to the assay, LCLs at 100,000 cells/well and MEFs at 60,000 cells/well. LCLs were treated with 200 nM rapamycin for 24 hours prior to running MitoStress Assay. To determine mitochondrial content, WT and S47 MEFs were incubated with 500 nM of MitoTracker Green (ThermoFisher Scientific, M7514) for one hour at 37°C. Cells were then spun down, washed once with PBS, spun down and resuspended in PBS. The FACSCelesta (BD Biosciences) was used to detect fluorescence and at least 10,000 events were measured per sample.

### Metabolite measurements

Media was collected after 24 hours after plating LCLs or MEFs, and the YSI-71000 Bioanalyzer was used to determine glucose, glutamine, lactate and glutamate levels as previously described (Londono Gentile et al., 2013). For the metabolic flux studies, cells were incubated in uniformly labeled ^13^C-glucose (25 mM) as indicated in the figure legends. For intracellular extracts, after incubation, the culture medium was aspirated and cells were washed once in ice-cold PBS. Metabolites were extracted by adding a solution of methanol/acetonitrile/water (5:3:2) to the well. Plates were incubated at 4°C for 5 minutes on a rocker and then the extraction solution was collected. The metabolite extract was cleared by centrifuging at 15,000 x *g* for 10 minutes at 4°C. Supernatants were transferred to LC-MS silanized glass vials with PTFE caps and either run immediately on the LC-MS or stored at -80°C. LC-MS analysis was performed on a Q Exactive Hybrid Quadrupole-Orbitrap HF-X MS (ThermoFisher Scientific) equipped with a HESI II probe and coupled to a Vanquish Horizon UHPLC system (ThermoFisher Scientific). 0.002 ml of sample is injected and separated by HILIC chromatography on a ZIC-pHILIC 2.1-mm. Samples were separated by ammonium carbonate, 0.1% ammonium hydroxide, pH 9.2, and mobile phase B is acetonitrile. The LC was run at a flow rate of 0.2 ml/min and the gradient used was as follows: 0 min, 85% B; 2 min, 85% B; 17 min, 20% B; 17.1 min, 85% B; and 26 min, 85% B. The column was maintained at 45°C and the mobile phase was also pre-heated at 45°C before flowing into the column. The relevant MS parameters were as listed: sheath gas, 40; auxiliary gas, 10; sweep gas, 1; auxiliary gas heater temperature, 350°C; spray voltage, 3.5 kV for the positive mode and 3.2 kV for the negative mode. Capillary temperature was set at 325°C, and funnel RF level at 40. Samples were analyzed in full MS scan with polarity switching at scan range 65 to 975 m/z; 120,000 resolution; automated gain control (AGC) target of 1E6; and maximum injection time (max IT) of 100 milliseconds. Identification and quantitation of metabolites was performed using an annotated compound library and TraceFinder 4.1 software. The “M+X” nomenclature refers to the isotopolog for that given metabolite. Isotopologs are chemically identical metabolites that differ only in their number of carbon-13 atoms. For instance, “M+2 citrate” means that two of the six carbons in citrate are carbon-13 while the other four are carbon-12. “M+4 citrate” means that four of the six carbons in citrate are carbon-13 while the other two are carbon-12.

### BMH crosslinking

Immortalized WT and S47 MEFs were generated and maintained as previously described (Jennis et al., 2016; Leu et al., 2019). For BMH crosslinking studies, the WT and S47 cells were cultured in 1% FBS DMEM medium and treated with PBS or 50 µM diethyl maleate (DEM, ThermoFisher Scientific AC114440010) for 5 h. Proteins were extracted from cultured cells or mouse tissue (skeletal muscle, lungs) using 1X DPBS (Thermo Fisher Scientific 14190144) supplemented with 0.5% IGEPAL CA-630, 1 mM PMSF, 6 µg/ml aprotinin, and 6 µg/ml leupeptin at 4°C. The tissues were homogenized using the Wheaton Overhead Stirrer. Total cellular homogenates were pulse sonicated using the Branson digital sonifier set at 39% amplitude. Total protein extracts (100 µg per reaction) were incubated with or without 1 mM BMH (Thermo Fisher Scientific 22330) for 30 min at 30°C. The samples were quenched with an equal volume of 2x Laemmli Sample Buffer (BioRad 1610737) supplemented with 5% β-Mercaptoethanol (BioRad 1610710) and heated for 10 min at 100°C. The protein samples were size fractionated on Novex 4-20% Tris-Glycine Mini Gels (Thermo Fisher Scientific XP04200BOX) at room temperature and subsequently transferred overnight onto Immuno-Blot PVDF membranes (BioRad 1620177) at 4°C. The membranes were blocked with 3% nonfat dry milk (BioRad 1706404) in 1X PBST for 30 min at room temperature and incubated with the GAPDH antibody (Cell Signaling Technology 2118) overnight with rotation/nutation at 4°C. After washing the blots in 1X PBST, the membranes were incubated with Donkey anti-Rabbit (Jackson ImmunoResearch 711-036-152) for 2 h at room temperature. Membrane-immobilized protein detection used ECL Western Blotting Detection Reagents (GE Healthcare RPN2106; Millipore Sigma GERPN2106).

### Proximity Ligation Assay

Cells were grown on Lab-Tek II 8-well chamber slides, were either untreated, treated with 50 µM diethyl maleate (DEM, ThermoFisher Scientific AC114440010) for 5 hours or treated with 10 µM of buthionine sulfoximine for 24 hours (BSO, Cayman Chemicals, 83730-53-4) and fixed with 4% paraformaldehyde (Electron Microscopy Sciences, 15710). Protein-protein interactions were assessed using the PLA Duolink in situ starter kit (Sigma Aldrich, DUO92101) following the manufacturer’s protocol. The following primary antibodies were used: Rheb 1:50 (B-12, Santa Cruz, sc271509), mTOR 1:500 (7C10, Cell Signaling, 2983), GAPDH 1:1000 (14C10, Cell Signaling, 2118). ImageJ software (NIH, Rockville, MD) was used to quantify PLA signals.

### Body composition and metabolic cage studies

WT and S47 mice in a pure C57Bl/6 background are previously described (Jennis et al., 2016). All mouse studies were performed in accordance with the guidelines in the Guide for the Care and Use of Laboratory Animals of the NIH and all protocols were approved by the Wistar Institute Institutional Animal Care and Use Committee (IACUC). Mice were fed an *ad libitum* diet and were housed in plastic cages with a 12-hour/12-hour light cycle at 22°C unless otherwise stated. Fat and lean content were measured in live male mice at 6 weeks of age using nuclear magnetic resonance (NMR) with the Minispec LF90 (Bruker Biospin, Billerica, MA). Indirect calorimetry was conducted to assess metabolic capabilities in mice (Oxyman/Comprehensive Laboratory Animal Monitoring System (CLAMS); Columbus Instruments). Six-week old mice were single caged, provided with water and food *ad libitum* and allowed to acclimate to the cages for 2 days. Oxygen consumption (VO_2_) and carbon dioxide production (VCO_2_) were recorded for 48 hours using an air flow of 600 ml/min and temperature of 22°C. Respiratory exchange ratio (RER) is calculated as VCO_2_/VO_2_ and heat (kcal/h) is calculated by 3.815 + 1.232*(RER). Photodetectors were used to measure physical activity (Optovarimex System; Columbus Instruments).

### Treadmill and serum metabolite studies

Mice were allowed to acclimate to the metabolic treadmill (Columbus Instruments) for 5 minutes before beginning their runs. The treadmill was then set to 5m/min and speed increased by 5m/min every 2 minutes until 20m/min was reached. Upon reaching 20 m/min, the incline was increased by 5 degrees every 2 minutes until reaching a maximum of 25 degrees. Mice were allowed to run at this maximum speed and incline until exhaustion, defined by the mice spending 10 continuous seconds on the shock grid. Lactate (Nova Biomedical) and glucose (One Touch) measurements were taken using test strips just prior to treadmill entry and immediately after exhaustion using handheld meters. Tail blood was also taken prior to treadmill entry and immediately after exhaustion and metabolites measured using the Vettest serum analyzer (Idexx Laboratories).

### Statistical Analysis

Unless otherwise stated, all experiments were performed in triplicate. The two-tailed unpaired Student t-test was performed. All *in vitro* data are reported as the mean ± SD unless stated otherwise, and *in vivo* are reported as the mean ± SE. Statistical analyses were performed using GraphPad Prism, p-values are as follows: (*) p-value < 0.05, (**) p-value < 0.01, (***) p-value < 0.001, (****) p-value < 0.0001. For the CLAMs and mouse exercise data, the Wilcoxon rank-sum test was used to compare the differences between S47 and WT mice.

## Acknowledgments

This work was supported by R01 CA102184 (M.E.M), R01 CA139319 (D.L.G. and M.E.M), P01 CA114046 (D.L.G. and M.E.M), F32 CA220972 (T.B.), R01 CA174761 (K.E.W.), R01 AG043483 (J.A.B.) and the Penn Diabetes Research Center (P30-DK19525). RSA is partly supported by a Bloomberg Distinguished Professorship. The authors would like to acknowledge the Histotechnology, Laboratory Animal and Imaging facilities at the Wistar Institute. The authors are grateful to Allie Lipshutz and Lindsey Schweitzer for technical help, and Matthew Jennis for the mouse weight data.

---

**Figure 1 – Supplement 1.**
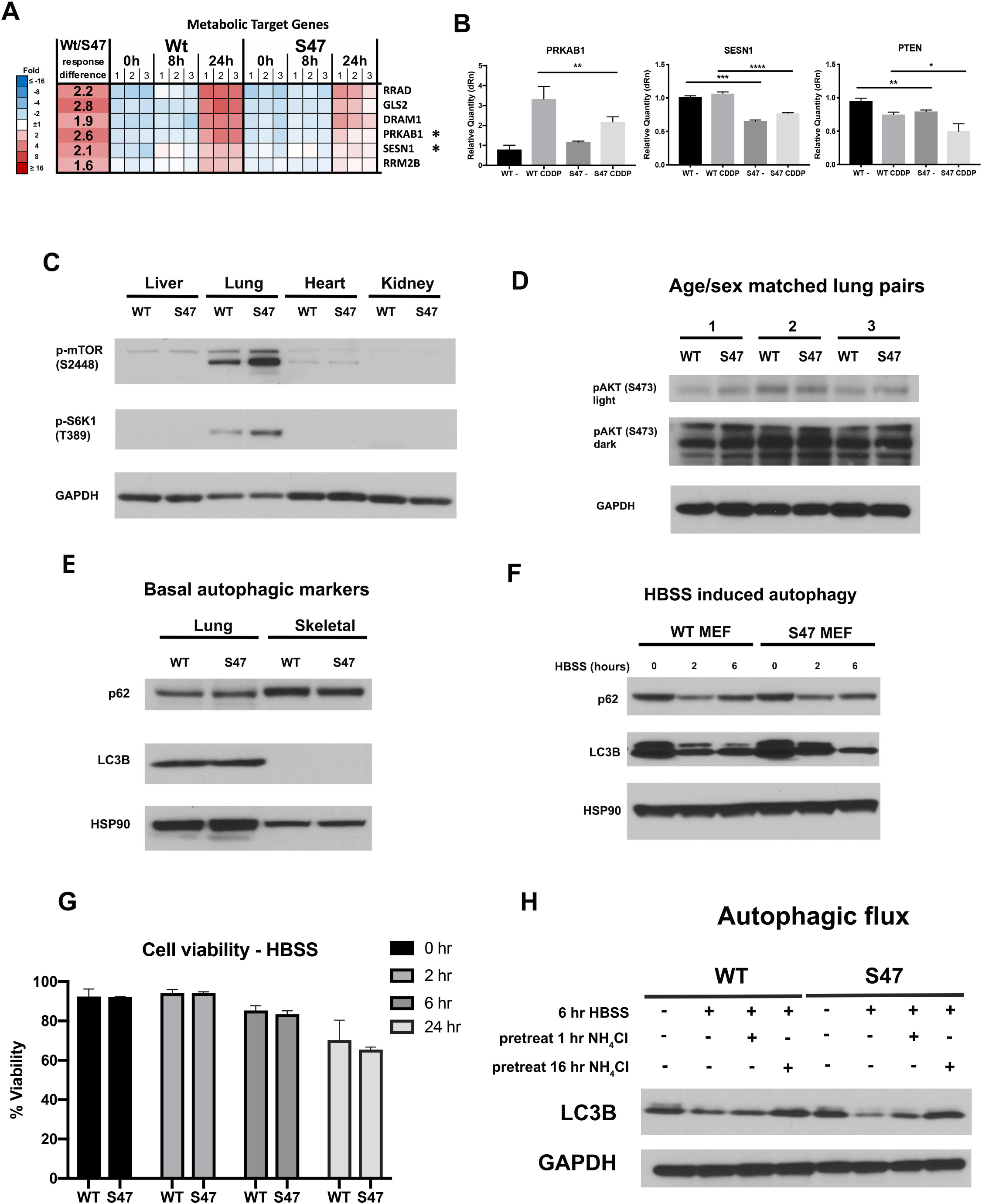
Altered metabolic markers in S47 cells and tissues. (A) Microarray analysis of WT and S47 LCLs treated with 10 µM cisplatin for 0, 8 & 24 hours reveal several metabolism genes as differentially expressed. (B) qRT-PCR analysis of *PRKAB1*, *SESN1* and *PTEN* in human LCLs treated with 10 µM cisplatin for 24 h. All values were normalized to a control gene (18S); n=4, error bars indicate standard deviation. (*) p-value < 0.05, (**) p-value < 0.01, (***) p-value < 0.001, (****) p-value < 0.0001, Student’s t-test. (C) Whole cell lysates were extracted from WT and S47 liver, lung, heart and kidney followed by Western Blot analysis for the proteins indicated. (D) Whole cell lysates were extracted from 3 WT and 3 S47 mouse lungs and analyzed by Western blot for the phospho-Akt and GAPDH (loading control). Light and dark exposures are shown. (E) Whole cell lysates were extracted from WT and S47 mouse lung and skeletal tissue and were subjected to Western blot analysis, probing for p62, LC3B and HSP90 (loading control). (F) WT and S47 MEFs were treated with HBSS for 0, 2 and 6 hours and were subjected to Western blot analysis for the indicated proteins. (G) WT and S47 MEFs were treated with HBSS for indicated time points and were subjected to viability analysis using Trypan Blue; n = 3, error bars indicate standard deviation. (H) Autophagic flux was measured in WT and S47 MEFs treated pretreated with NH_4_Cl for indicate time points, followed by HBSS treatment for 6 hours. Cell lysates were subjected to Western blot analysis and immunoblotted for LC3B and GAPDH (loading control).

**Figure 2 – Supplement 1.**
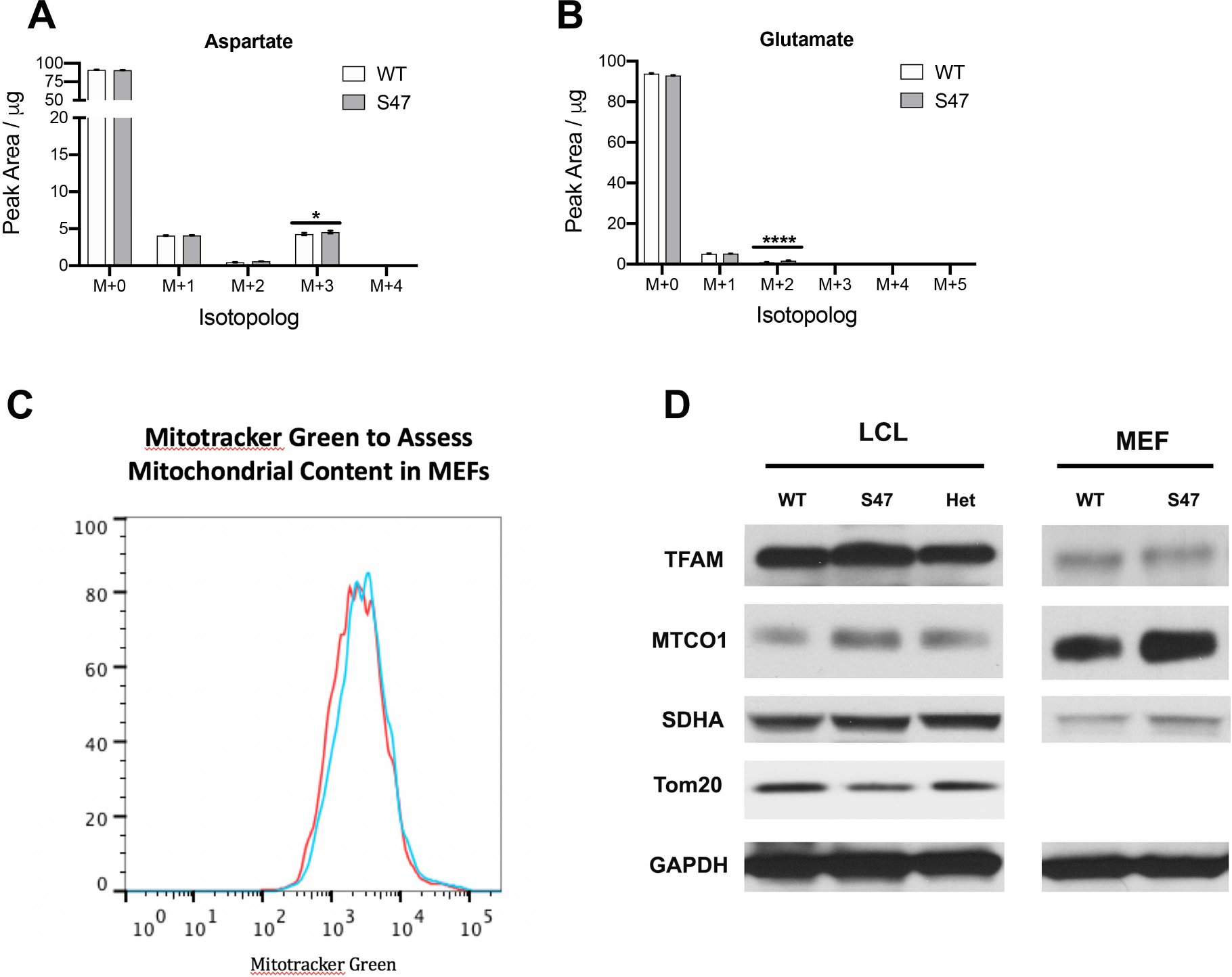
No obvious change in mitochondrial content in S47 cells. (A-B) WT and S47 MEFs were incubated with 25 mM ^13^C-glucose for 15 minutes and the abundance of aspartate (A) and glutamate (B) isotopologs was quantified by LC-MS/MS. Data are presented as mean ± SD, n =3. (C) Mitochondrial mass in WT and S47 MEFs measured by Mitotracker Green fluorescence. Data depicted are representative of three independent experiments performed in triplicate. (D) Cell lysates extracted from WT and S47 LCLs and MEFs were subjected to Western blot analysis and immunoblotted for TFAM, MTCO1, SDHA, TOMM20 and GAPDH (loading control).

**Figure 4 – Supplement 1.**
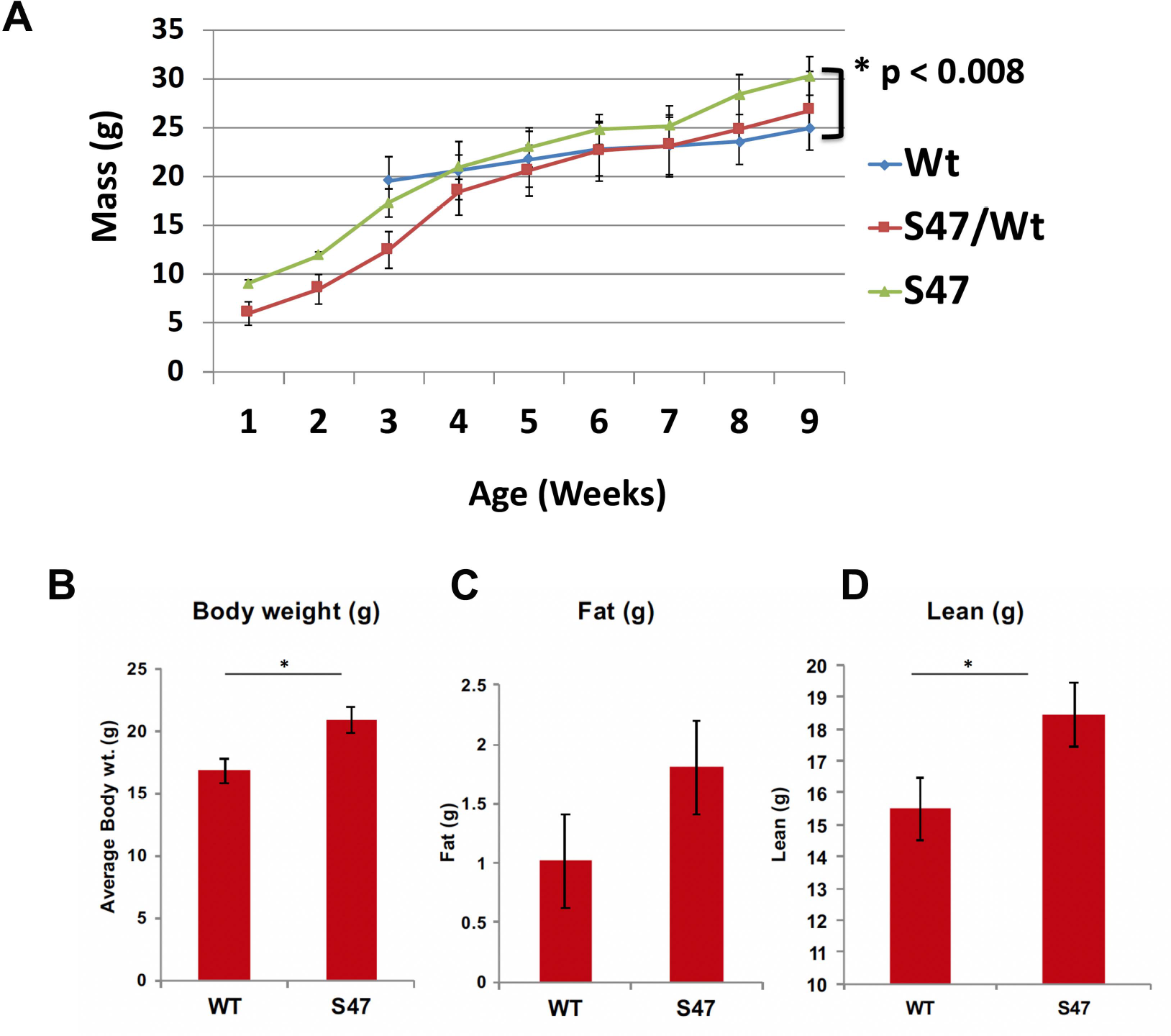
Increased size in S47 mice compared to WT littermates. (A) Progression of mouse weight in WT, S47 and heterozygous WT/S47 mice over the course of 18 weeks; a minimum of 10 mice of each genotype were tracked, and error bars mark standard error. (B-D) Body weight, fat content and lean content as measured by proton magnetic resonance spectroscopy (H-MRS); n=5 mice per genotype, error bars mark standard error, (*) p-value < 0.05.

**Figure 5 – Supplement 1.**
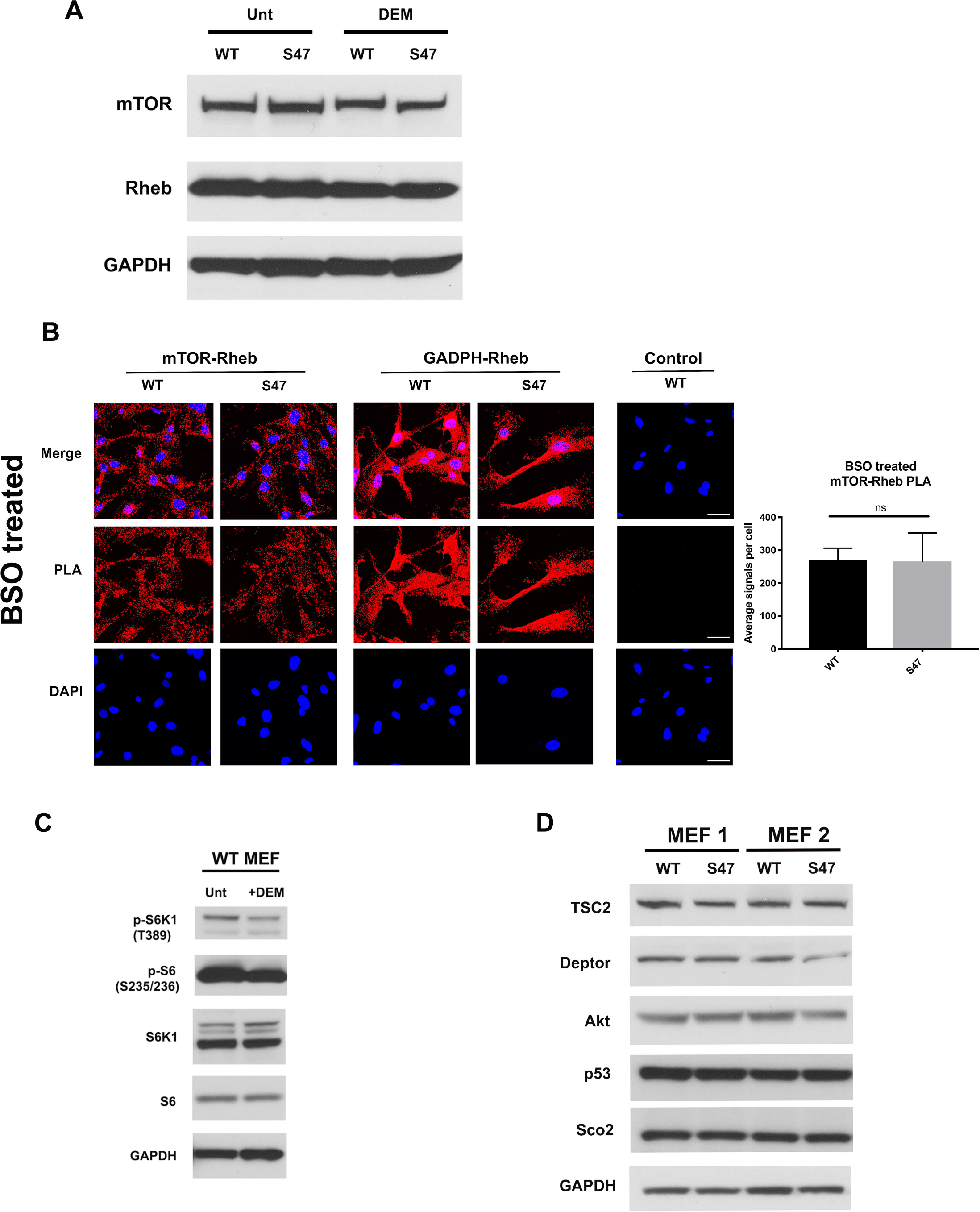
Impact of glutathione depletion in S47 cells. (A) WT and S47 MEFs were untreated or treated with 50 µM of DEM for 5 h. Cell lysates were subjected to Western blot analysis and immunoblotted for total mTOR, Rheb and GAPDH. (B) An *in situ* proximity ligation assay was performed in WT and S47 MEFs that are treated with 10 µM of BSO for 24 h. Shown on the right is the quantification of the mTOR-Rheb interactions, measured as the average number of PLA signals per nuclei. (C) Cell lysates were extracted from WT MEFs treated with 50 µM of DEM for 5 h. Cell lysates were subjected to Western blot analysis and immunoblotted for the proteins indicated. (D) Cell lysates were extracted from two sets of WT and S47 MEFs and were analyzed by Western blot for TSC2, DEPTOR, AKT, p53, Sco2 and GAPDH.

**Figure 6 – Supplement 1.**
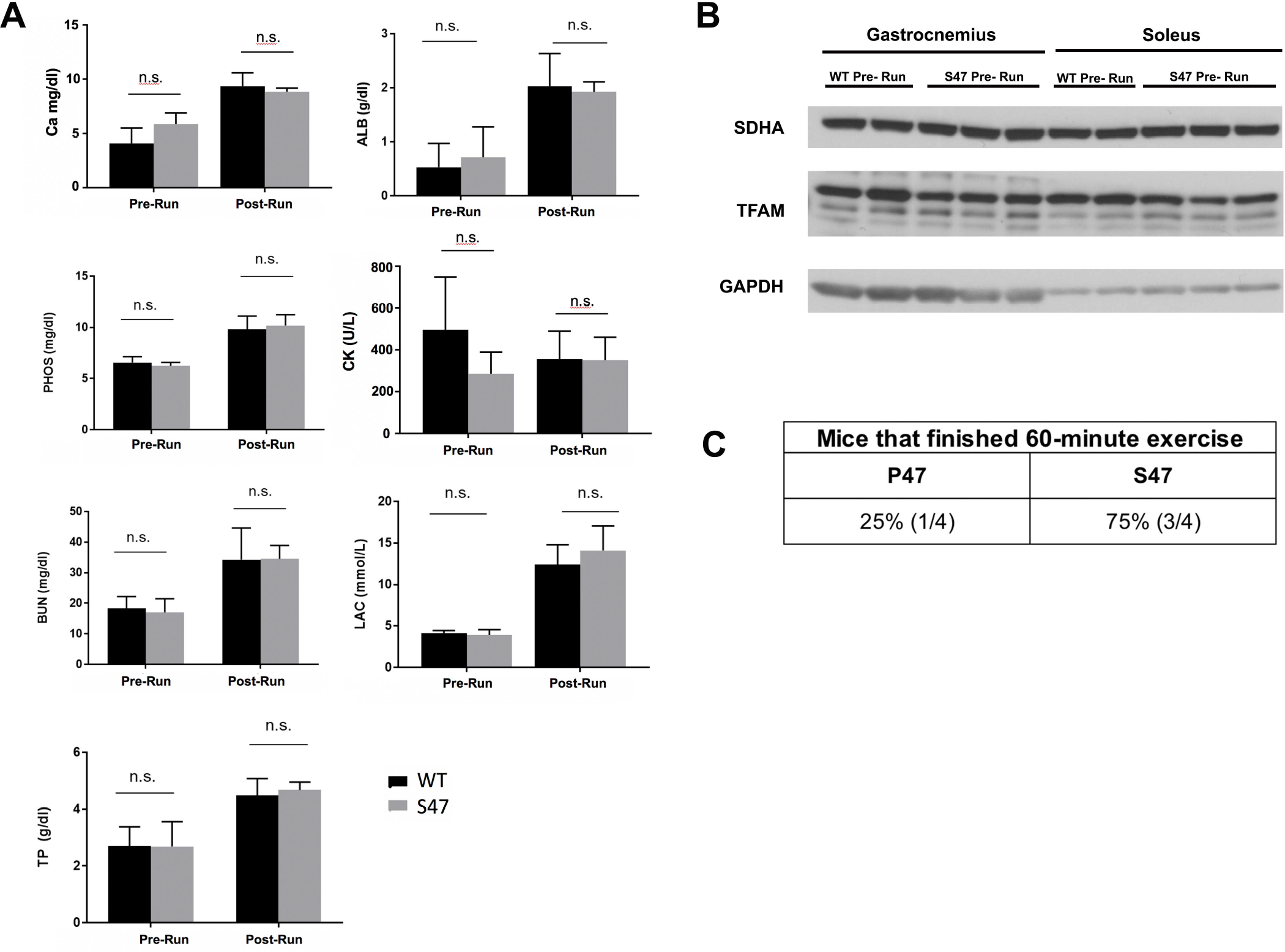
Serum metabolites and protein markers pre- and post-exercise. (A) The following blood serum metabolite levels measured in WT and S47 mice before and after treadmill run: Ca (calcium), ALB (albumin), PHOS (phosphate), CK (creatine kinase), BUN (blood urea nitrogen), LAC (lupus anticoagulant), TP (total protein). n.s. not significant, Student’s t-test. (B) Whole cell lysates were extracted from gastrocnemius or soleus muscle from untreated WT and S47 mice and subjected to Western Blot analysis for the proteins indicated. (C) WT and S47 mice were subjected to a 60 minute treadmill run. Table indicates proportion of mice that completed run; n = 4.

